# Cytokine expression patterns: A single-cell RNA sequencing and machine learning based roadmap for cancer classification

**DOI:** 10.1101/2023.06.01.542694

**Authors:** Zhixiang Ren, Yiming Ren, Pengfei Liu, Huan Xu

**Affiliations:** Peng Cheng Laboratory, Shenzhen, Guangdong Province, China 518055; Shanghai Frontiers Science Center of Optogenetic Techniques for Cell Metabolism and Shanghai Key Laboratory of New Drug Design, School of Pharmacy, East China University of Science and Technology, Shanghai, China 200237; School of Computer Science and Engineering, Sun Yatsen University, Guangzhou, Guangdong Province, China 528406

**Keywords:** Tumor immune microenvironment, Cancer classification, Machine learning, Single-cell RNA sequencing

## Abstract

Cytokines are small protein molecules that exhibit potent immunoregulatory properties, which are known as the essential components of the tumor immune microenvironment (TIME). While some cytokines are known to be universally upregulated in TIME, the unique cytokine expression patterns have not been fully resolved in specific types of cancers. To address this challenge, we develop a TIME single-cell RNA sequencing (scRNA-seq) dataset, which is designed to study cytokine expression patterns for precise cancer classification. The dataset, including 39 cancers, is constructed by integrating 695 tumor scRNA-seq samples from multiple public repositories. After screening and processing, the dataset retains only the expression data of immune cells. With a machine learning classification model, unique cytokine expression patterns are identified for various cancer categories and pioneering applied to cancer classification with an accuracy rate of 78.01%. Our method will not only boost the understanding of cancer-type-specific immune modulations in TIME but also serve as a crucial reference for future diagnostic and therapeutic research in cancer immunity.

## I. Introduction

**B**ESIDES malignant cells, there are many other components in tumors, including stroma cells, immune cells, hormones, cytokines and many other biologically-active molecules, forming a well-organized complex system [1]. TIME consists of the immunological components within tumors, which is known to be associated with tumor development, progression, metastasis and immune escape [2]. Immune cells in TIME, including monocytes, macrophages, neutrophils, dendritic cells (DCs), natural killer (NK) cells, lymphocytes, and rare progenitors like common myeloid progenitors (CMPs) or hematopoietic stem cells (HSCs), interact with each other and tumor cells through cell signaling molecules, constituting a complicated regulatory network in TIME [3], [4]. Identifying the expression patterns of these signaling molecules not only reveals immune cell regulations in TIME but also provides essential information for the modeling of the TIME immune network.

Cytokines are small but essential proteins that exert potent immunoregulatory roles in physiological and pathological conditions. More than 200 cytokines have been identified so far, which can be categorized into different families, including interleukins (ILs), colony stimulating factors (CSFs), interferons (IFNs), tumor necrosis factors (TNFs), growth factor (GFs) and chemokines [5]. Considered as the hormones of the immune system, many cytokines, such as TNF-*α*, IL-1*β*, IL-2, IL-6, IL-12 and IL-15, have already been utilized in cancer immunotherapies either to enhance growth inhibitory effects or suppress tumor-promoting actions [6], [7]. While a histology-independent uniform cytokine cascade is noticed in different types of cancers, unique cytokine expression patterns are also being identified in specific cancer types [8], [9], which may be developed as potential diagnostic biomarkers or therapeutic targets [6], [10], [11].

Recent advances in single-cell RNA sequencing (scRNAseq) technique have enabled us to study gene expressions at the RNA level in individual cells isolated from tumor tissues [12]. The scRNA-seq data deposited in various public repositories can be utilized for comprehensive analysis of TIME. For example, CancerSCEM launched an online analytical platform of TIME based on the scRNA-seq data of 208 cancer samples from 28 projects [13]. TMExplorer created an R package for gene expression analysis in the 48 processed datasets representing 26 different human cancer types and 4 different mouse cancer types [14]. Although these studies provided useful tools for the visualization of TIME data, the outputs are too general to reflect the specific cytokine expressions in individual cancer types.

Thanks to the advancement of artificial intelligence technology, several methods that combine scRNA-seq data and machine learning (ML) have been developed for precise cancer classification. [15] proposed a neural network-based classifier that differentiates between 21 types of cancer and normal tissues using scRNA-seq data, achieving high accuracy. [16] utilized gene expression of scRNA-seq data to predict the relationship with somatic mutation and used ML methods for pan-cancer classification. Additionally, a method proposed in [17] extracted effective genes from RNA-seq data, thereby providing performance gains for a pan-cancer classifier based on scRNA-seq data. Although these methods have investigated gene expression patterns in tumor scRNA-seq data, the immunoregulatory mechanism of cytokines for specific cancer types has not been revealed.

In this paper, in order to identify the precise cytokine expression patterns in TIME of different cancers, scRNA-seq data for 39 types of cancers is collected from multiple public repositories, which includes 695 immune cell-containing tumor samples. A carefully designed workflow, as shown in Figure 1, is employed for data processing. Initially, predefined marker genes are used to screen the TIME. Furthermore, the R library SingleR is utilized to annotate immune cell types in TIME. To improve the accuracy of classification, we replace the specific genes obtained by SingleR with validated marker genes. After the visualization of cancer-type-specific immune cell-cytokine relations, the ML classification model is employed to identify the cytokine expression patterns, achieving a cancer classification accuracy of 78.01%. So far, this is the very first study focusing on unique cytokine expression patterns in TIME for classifying various types of cancers.

**Fig. 1.**
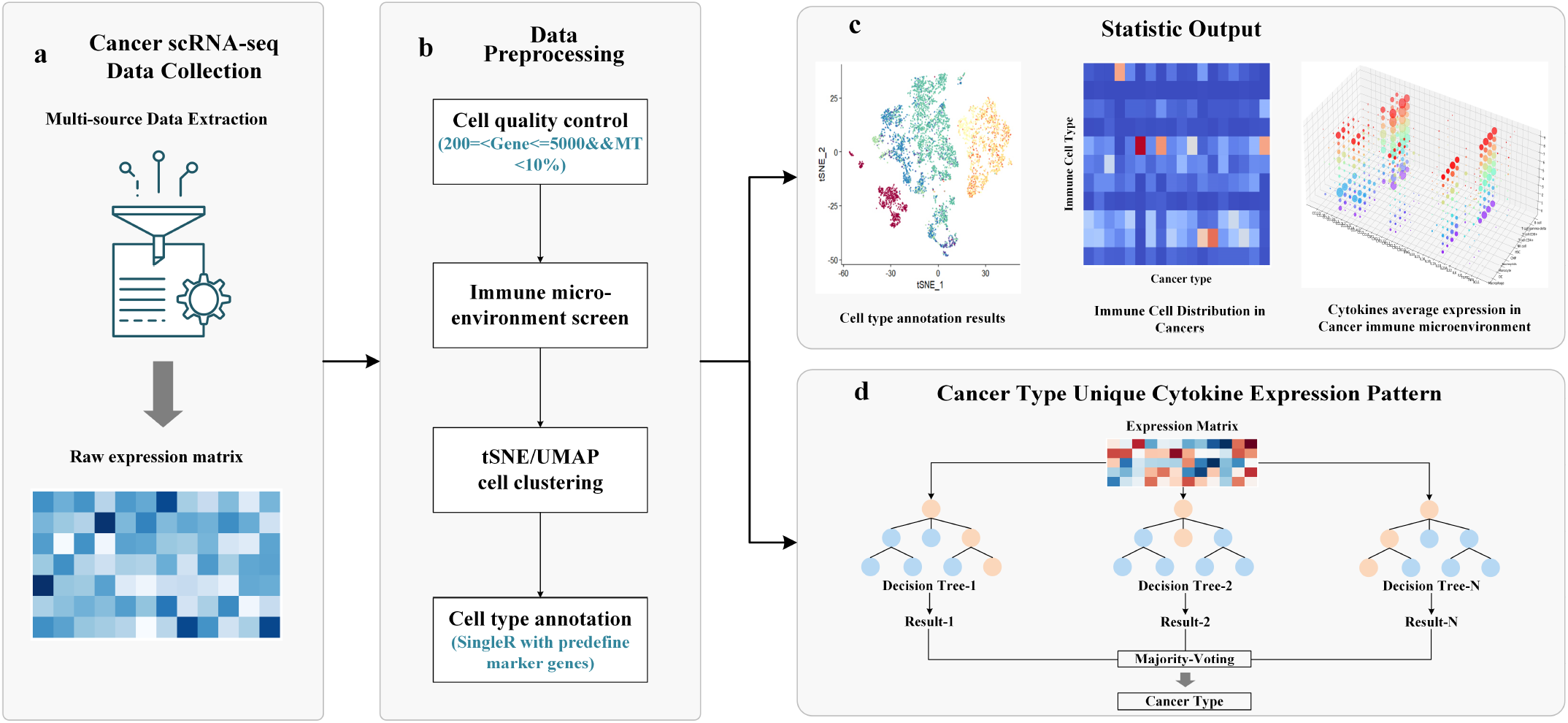
Workflow of Data Processing and Analysis. **a.Cancer scRNA-seq Data Collection** devises a custom Python script to effectively gather the raw expression matrix of scRNA-seq data pertaining to the TIME from a diverse range of cancer patients, primarily drawing from public databases, such as GEO. **b. Data Preprocessing** utilizes the Seurat library for cell quality control(200 ≤gene ≤5000 and MT***<***10%). The validated marker genes are then used for TIME screening and as the template for SingleR immune cell type annotation. **c. Statistic Output** shows various ways to visualize the general cytokine expression patterns in the TIME of various cancer types, including tSNE,uMAP clustering. **d. Cancer Type Unique Cytokine Expression Pattern** models the correlation between cytokine expression data of the TIME and specific cancer types with the Random Forest classifier.

## II. METHODS

### A. Cancer scRNA-seq data collection and processing

The dataset used in this study is collected from multiple public scRNA-seq databases, including GEO (https://www.ncbi.nlm.nih.gov/geo/) [18], ArrayExpress (https://www.ebi.ac.uk/biostudies/arrayexpress) [19], Single Cell Expression Atlas (https://www.ebi.ac.uk/gxa/sc/home) [20], and ZENODO (https://zenodo.org/). A total of 695 human tumor tissue samples are collected and processed from 117 scRNA-seq projects covering 39 human cancers in Table S1. The expression matrix of raw count and R package Seurat v4.3.0 (https://satijalab.org/seurat/) are utilized for single-cell analysis workflow, including cell quality control (200 ≤ nfeatures ≤ 5000 and MT *<* 10%), PCA (principal component analysis) dimension reduction, UMAP (uniform manifold approximation and projection) and tSNE (t-distributed stochastic neighbor embedding) clustering with specific parameters and resolutions [21]. After removing low-quality cells, the TIME is initially screened based on marker genes. According to the marker genes on CancerSCEM documentation [13], 7 immune cell types, including B cell, GMP, HSC, Macro/Mono/DC, Neutrophil, NK cell, T cell, are defined to comprise the TIME of this study. The marker genes utilized for immune cell identification in TIME are shown in Table S2. The processed datasets can be accessed on ZENODO (https://sandbox.zenodo.org/record/1180710)

### B. Immune cell annotation in TIME

Currently, other than manual annotation, there is no standard method for immune cell identification in single-cell analysis. However, manual operation based on a couple of literature is susceptible to subjectivity. SingleR (single cell recognition) is a novel computational framework for cell annotation of scRNA-seq data by reference to aggregate transcriptomes, which has become one of the most trusted tools for cell type recognition in recent years [22]. To determine the diverse immune cell types in TIME, the cell type annotation is implemented with SingleR v2.0.0 (https://github.com/dviraran/SingleR). A total of 11 types of human immune cells are focused in this study, which are B cell, CMP, HSC, DC, Monocyte, Macrophage, Neutrophil, NK cell, T cell: CD4+, T cell: CD8+ and T cell: gamma-delta (Figure S1). Human Primary Cell Atlas (HPCA) [23] consists of 713 microarray samples derived from 37 primary cell types and 157 subtypes of human cells, including all the cell types of interest in TIME. Therefore, HPCA is adopted as the reference dataset to assist in cell type recognition, and only 368 samples contained the 11 cell types of interest are retained. In addition, to improve the accuracy of immune cell recognition, a set of typical marker genes for the concerned cell types in TIME are customized according to previous studies [22], [24], [25]. The customed marker genes are shown in Table I, which are used to replace the differential expression genes between samples as determined by SingleR when calculating Spearman correlation. The minimum difference median (*>* 0.1), which quantifies the difference between the assigned label and all other labels, is used to filter the unreliable annotation. Figure S1 represents the recognition effect of SingleR, and the average error rate is less than 5% across all samples.

**TABLE I.**
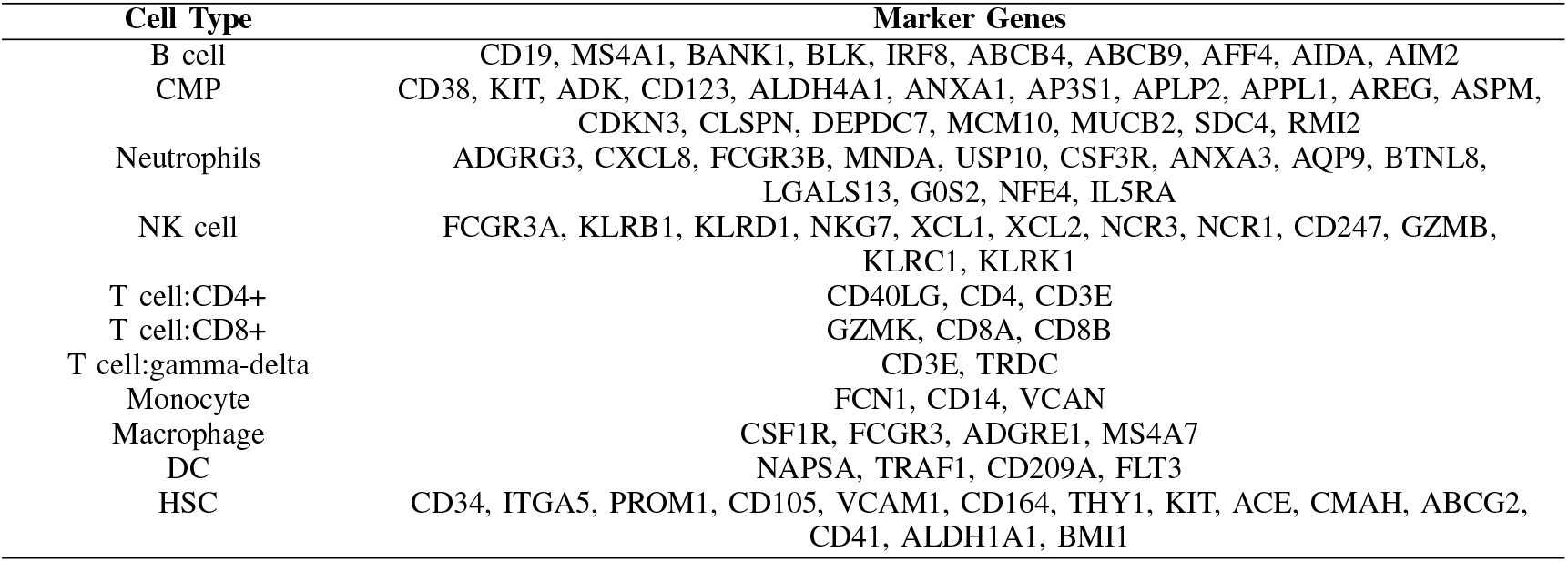
Marker genes for cell type recognition.

| Cell Type | Marker Genes |
| --- | --- |
| B cell | CD19, MS4A1, BANK1, BLK, IRF8, ABCB4, ABCB9, AFF4, AIDA, AIM2 |
| CMP | CD38, KIT, ADK, CD123, ALDH4A1, ANXA1, AP3S1, APLP2, APPL1, AREG, ASPM, CDKN3, CLSPN, DEPDC7, MCM10, MUCB2, SDC4, RMI2 |
| Neutrophils | ADGRG3, CXCL8, FCGR3B, MNDA, USP10, CSF3R, ANXA3, AQP9, BTNL8, LGALS13, G0S2, NFE4, IL5RA |
| NK cell | FCGR3A, KLRB1, KLRD1, NKG7, XCL1, XCL2, NCR3, NCR1, CD247, GZMB, KLRC1, KLRK1 |
| T cell:CD4+ | CD40LG, CD4, CD3E |
| T cell:CD8+ | GZMK, CD8A, CD8B |
| T cell:gamma-delta | CD3E, TRDC |
| Monocyte | FCN1, CD14, VCAN |
| Macrophage | CSF1R, FCGR3, ADGRE1, MS4A7 |
| DC | NAPSA, TRAF1, CD209A, FLT3 |
| HSC | CD34, ITGA5, PROM1, CD105, VCAM1, CD164, THY1, KIT, ACE, CMAH, ABCG2, CD41, ALDH1A1, BMI1 |

### C. Identification of cancer type-unique cytokine expressions in TIME with ML

To investigate the specific cytokine expression patterns in various types of cancers, we employed the constructed dataset in conjunction with ML. Random forest is applied for cytokine identification with the Python package Scikit-learn [26], and F1 score is introduced to evaluate the classification accuracy of each category. Since variations in the quantity of cells in each processed scRNA-seq sample may result in classification bias, the expression matrix of each sample is converted into a 106×11 matrix by averaging the expression of 106 cytokines across the 11 immune cell types, which ensured the uniformity of characteristics and eliminated the cell number differences between samples. In addition, as shown in Figure 2, the proportion of immune cells in each sample is appended to the final dimension of the training data, as the ratio of immune cells is regarded as an important feature for cancer classification. Given the substantial variance in the amount of samples across 39 cancer categories, the 10 cancers with the largest sample quantity are selected to improve the generalizability of the ML model.

**Fig. 2.**
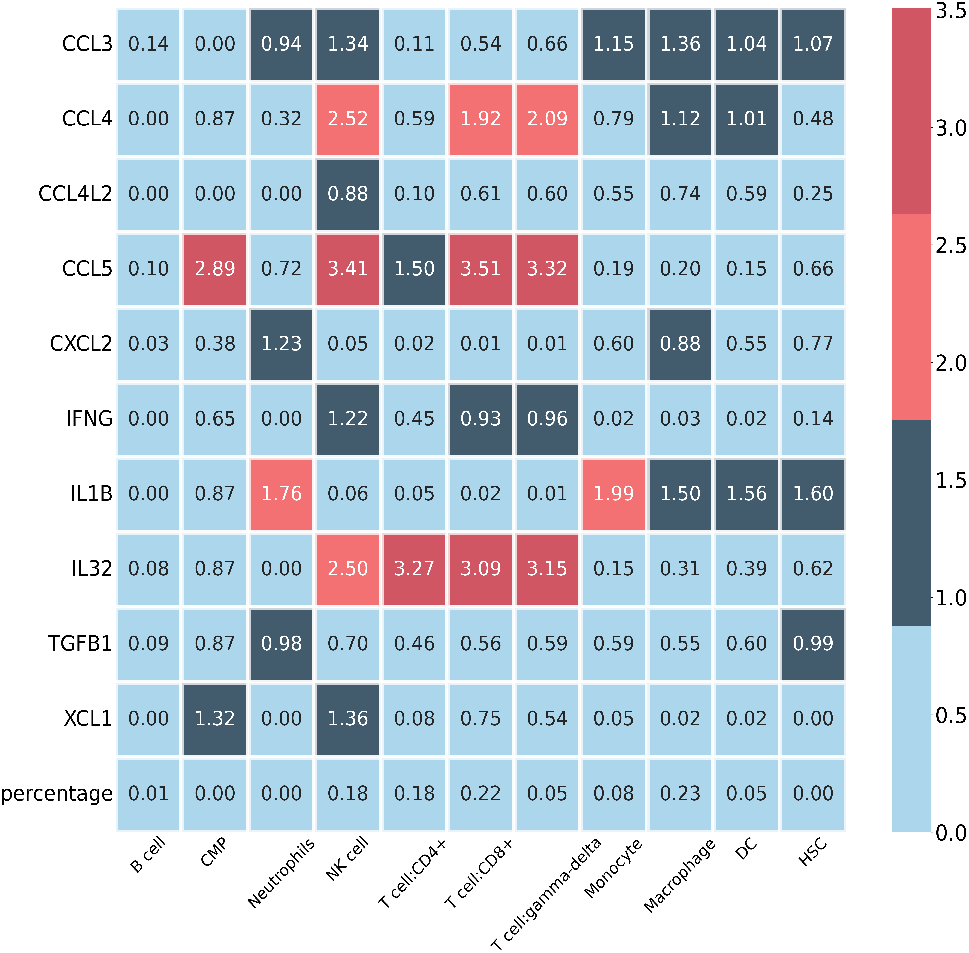
Format of Training Data. This figure shows the format of training data which contains the average expression of selected cytokines in specific immune cell types and immune cell type proportion in the sample. The rows contain two types of information: the selected cytokines and the proportion of each type of immune cell in the sample. The columns represent the 11 types of immune cells involved in the study.

## III. RESULTS

### A. Data statistics and visualization of TIME

After the cell type recognition with SingleR, tSNE and UMAP are applied to visualize the distribution of cells in the TIME (Figure 3 A and B). In order to investigate cytokine expressions in TIME, highly expressed cytokines in various types of immune cells (percentage*>*10%) in each sample are listed in a bubble chart Figure 3C. A total of 18 cancer categories are chosen for further analysis based on their sample sizes (amount of samples ≥10). The immune cell types in the TIME of the selected cancer samples and the top 20 expressed cytokines in the 18 types of cancers are also demonstrated in Figure S2. In addition, the correlations between cytokine expressions and immune cell types in the 10 cancer categories with the biggest sample sizes are summarized in a three-dimensional graph, visualizing the general cytokine expression patterns in the TIME of various cancer types (Figure 4). The average expressions of cytokines across all cancer types are also analyzed, identifying the commonly expressed cytokines in TIME, such as CCL4, IL32, CCL5, CCL3, CCL4L2, IL1B, TGFB1, CCL3L1, CXCL2, and TNFA, which are being ruled out for the recognition of unique cytokines for certain cancer types (Figure 5).

**Fig. 3.**
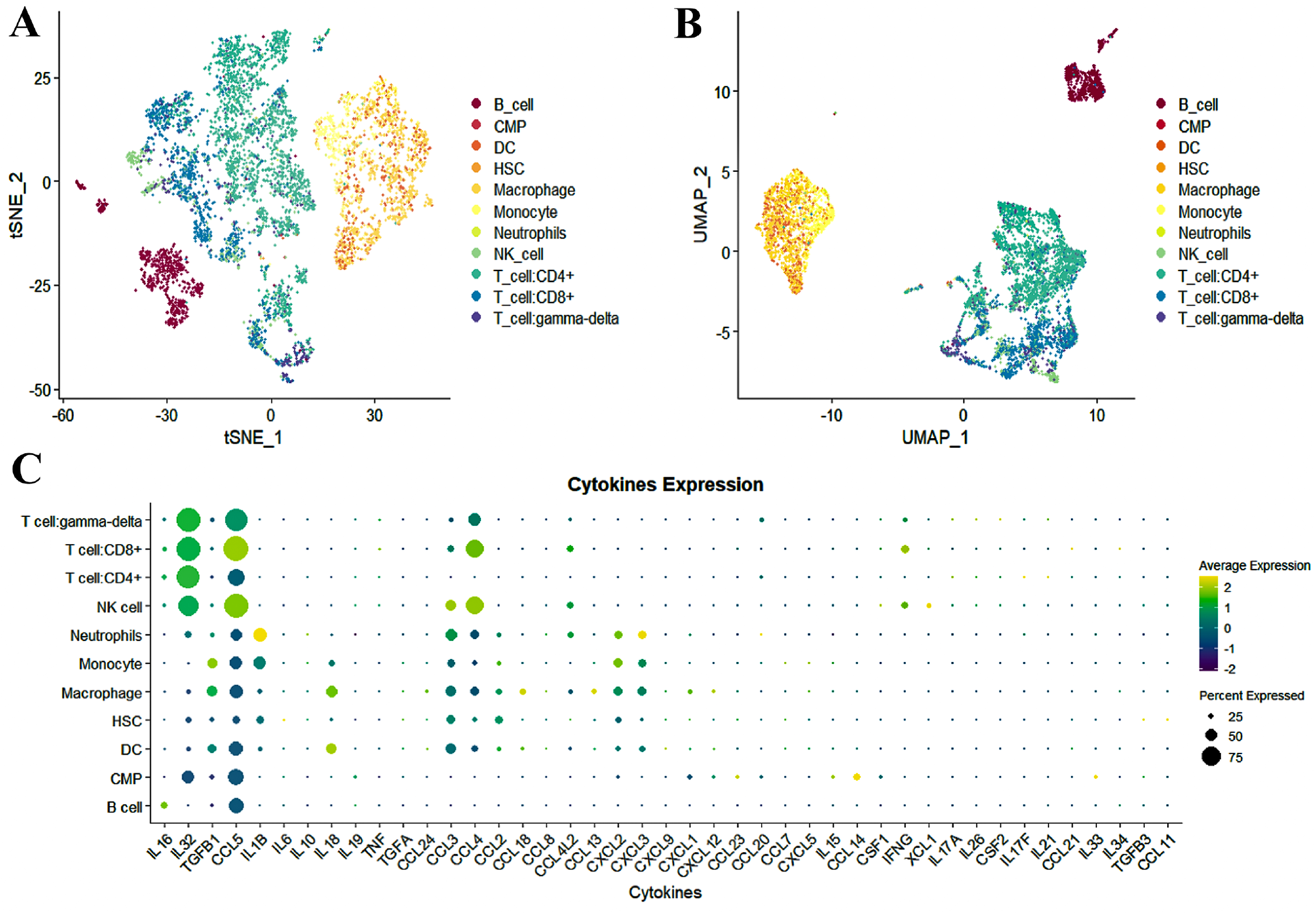
Showcase of Immune Cell Cluster and Cytokine Expression. **A, B. tSNE and UMAP Clustering Results**, represent the distribution of different types of immune cells in the microenvironment. **C. Cytokine Expression in Specific TIME** shows the average expression of cytokines in each immune cell type. Cytokines with an expression percentage lower than 10% are filtered out.

**Fig. 4.**
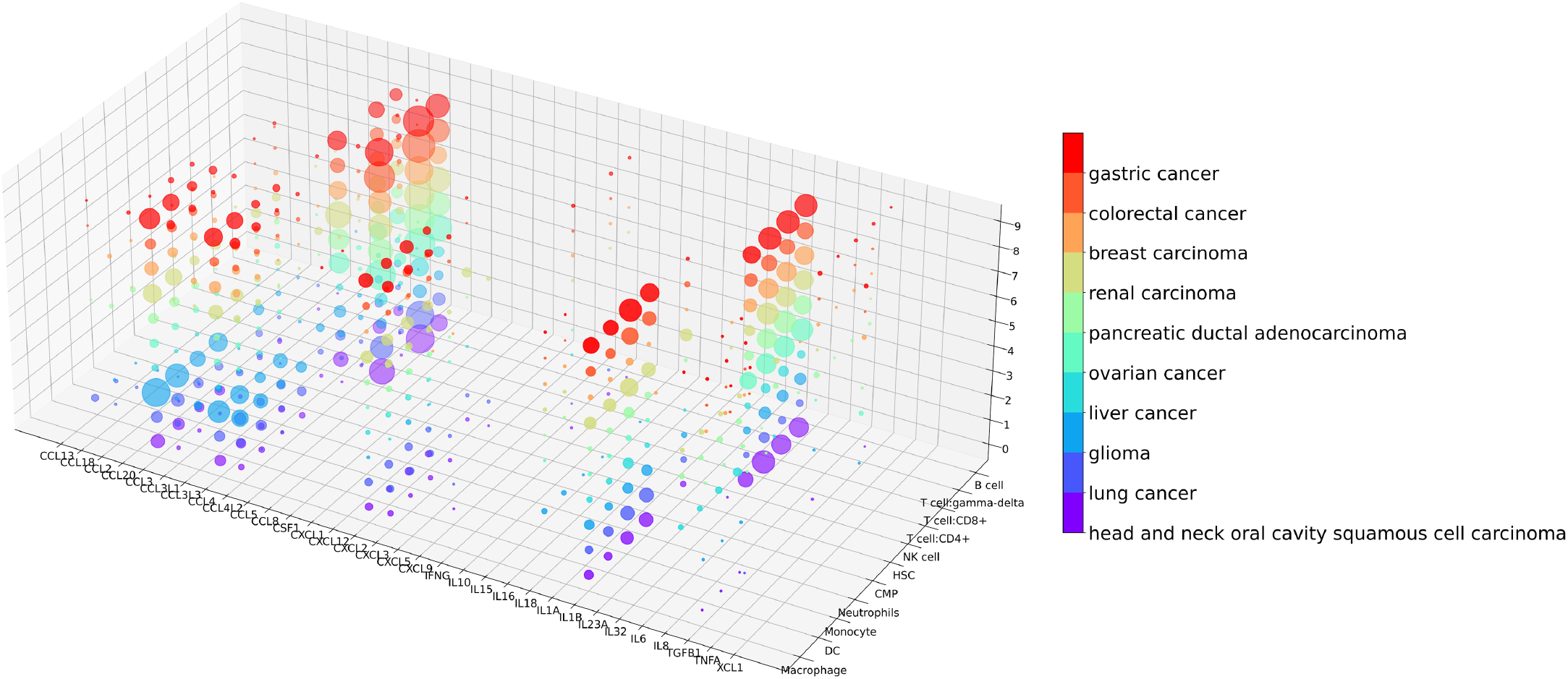
Top Cytokines Expression Patterns in TIME. Three-dimensional graph represents the expression of the top 20 cytokines in immune cells of the 10 cancers with the most samples.

**Fig. 5.**
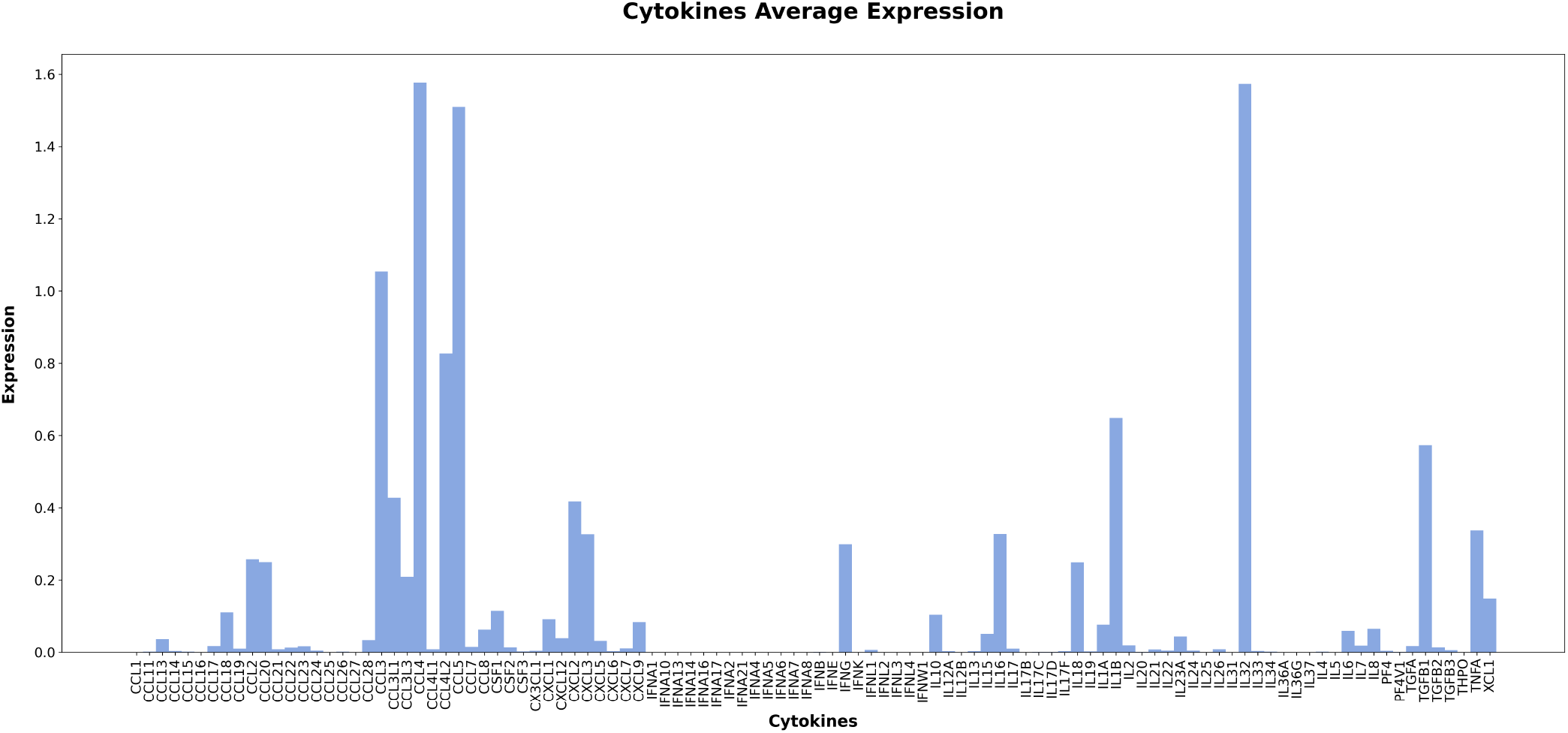
Global Cytokine Expression Pattern. The figure shows the distribution of cytokines average expression across all samples, with the top 10 highly-expressed cytokines being CCL4, IL32, CCL5, CCL3, CCL4L2, IL1B, TGFB1, CCL3L1, CXCL2, and TNFA.

### B. Identification of specific cytokines with ML

After the performance evaluation of decision tree, random forest, and multi-layer perceptron (MLP) models (Figure S3), random forest is applied in this study to verify the identification efficiency of the model in 3 to 10 types of cancers. The results of random forest classifying with 106 cytokines are shown in Table II and Figure S4, which demonstrated that the accuracy of the top three cancer types with the most samples could achieve a high precision of 92%. Although the accuracy declined progressively as the number of cancer samples decreased, the classification re-sults of 10 cancer types still achieved a remarkable performance of nearly 80%. Moreover, to explore highly-distinct cytokine expressions associated with each cancer type, the specific cytokines are extracted using the FindMarkers function of the scran v1.26.1 (https://bioconductor.org/packages/release/bioc/html/scran.html). The generated matrix is described as the concatenation of multiple samples corresponding to each cancer type. By continuously adjusting the threshold, we are able to identify multiple groups of specifically-expressed cytokines. In the threshold selection part, only the top 10 differential genes with a threshold (p-value *<* 0.01) are considered due to the high differential expression. To ensure that at least one specific cytokine is identified for each type of cancer, the minimum number of specific genes is set at 41. In addition, 65 is set as the upper limit, since the model effect diminished when the number of specific genes exceeded 65. The identified unique cytokines in each cancer type with 41 or 65 differential genes are listed in Table III and Table IV. The cytokines are obtained based on Scran’s FindMarkers function with a relatively strict threshold. Only the top 10 genes with p-value *<* 0.01 are selected. The application with 41 cytokines resulted in a reduction in classification accuracy but still had a minimum of 71.5%. However, as the number of selected cytokines increased, the model’s accuracy significantly dropped, indicating that the number of cytokines should be limited to an appropriate range in this model (Figure S4). In addition, the accuracy of ovarian cancer is consistently low, and the F1-score decreased considerably from 0.69 to 0.57 when specific cytokines are used, suggesting the complexity of cytokine expressions in the particular cancer type. For other cancer types, the identified cytokines with the lowest p-values are consistent with threshold alterations, which are summarized in Figure 6. These results will not only benefit the understanding of immune cell regulations in TIME but also provide valuable information for cancer immunity-related diagnostic and therapeutic research in the future.

**TABLE II.** Classification accuracy of random forest with different cytokines.

| Number of categories | F1@106 | F1@41 | F1@61 | F1@65 |
| --- | --- | --- | --- | --- |
| 3 | 92.63% | 90.52% | 90.95% | 91.81% |
| 4 | 89.14% | 86.06% | 88.14% | 87.45% |
| 5 | 86.31% | 81.94% | 84.88% | 85.2% |
| 6 | 83.23% | 77.34% | 78.13% | 79.95% |
| 7 | 80.66% | 74.06% | 78.63% | 79.33% |
| 8 | 78.81% | 74.17% | 75.9% | 77% |
| 9 | 77.61% | 73.45% | 74.88% | 76.31% |
| 10 | 78.01% | 71.55% | 75.62% | 75.81% |

**TABLE III.** 41 differential cytokines of top 10 cancer types.

| Cancer Type | Differential Cytokines |
| --- | --- |
| breast carcinoma | IL21, CCL19, CCL3L3, CXCL9, CCL8, IFNL4, IL17B |
| head and neck oral cavity squamous cell carcinoma | CCL22, CCL21, IL17F |
| lung cancer | CCL18, CCL13, CCL23, CXCL5, IL24, CCL17, IL21, IL37, CCL8 |
| gastric cancer | CCL24, IL17F, IL17D |
| colorectal cancer | IL17, IL22, IL17F, CCL15, IL26 |
| ovarian cancer | XCL1 |
| renal carcinoma | CCL14, CX3CL1 |
| glioma | CCL3L1, CCL2, IL1A, IL18, IL1B, IL10, IL6, CCL8, CCL4L2, CCL20 |
| pancreatic ductal adenocarcinoma | IL8, CCL18, CCL4L1 |
| liver cancer | PF4, CXCL7, CCL16, PF4V1, CCL15, THPO, IL8 |

**TABLE IV.**
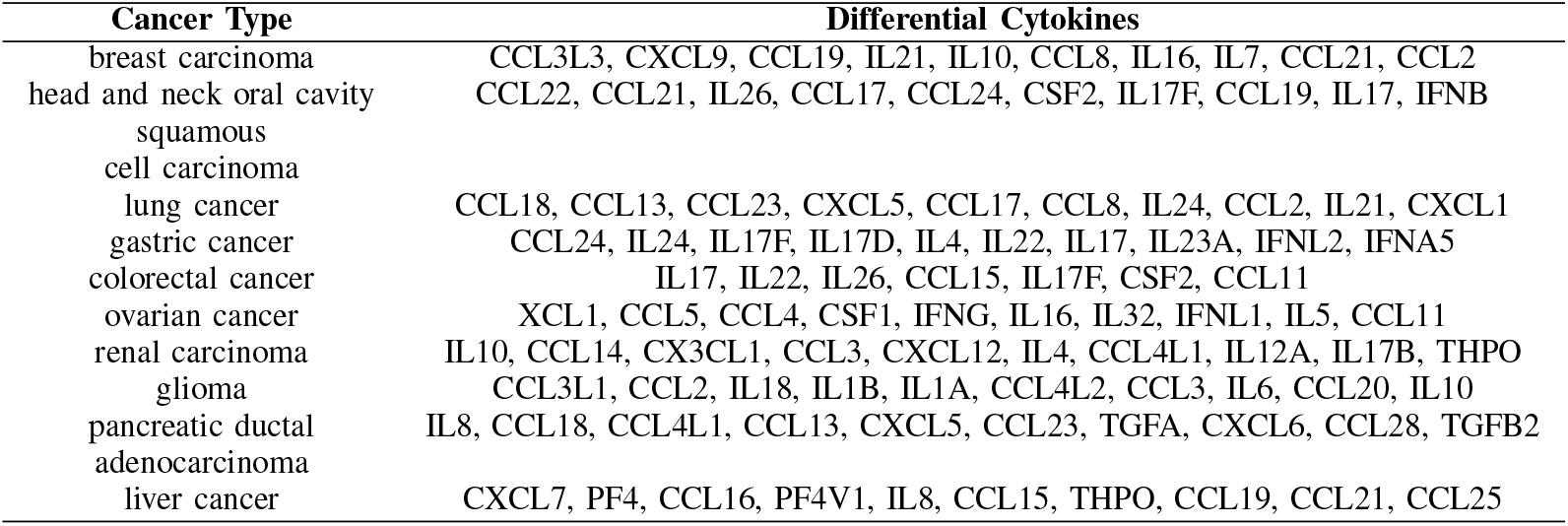
65 differential cytokines of top 10 cancer types.

**Fig. 6.**
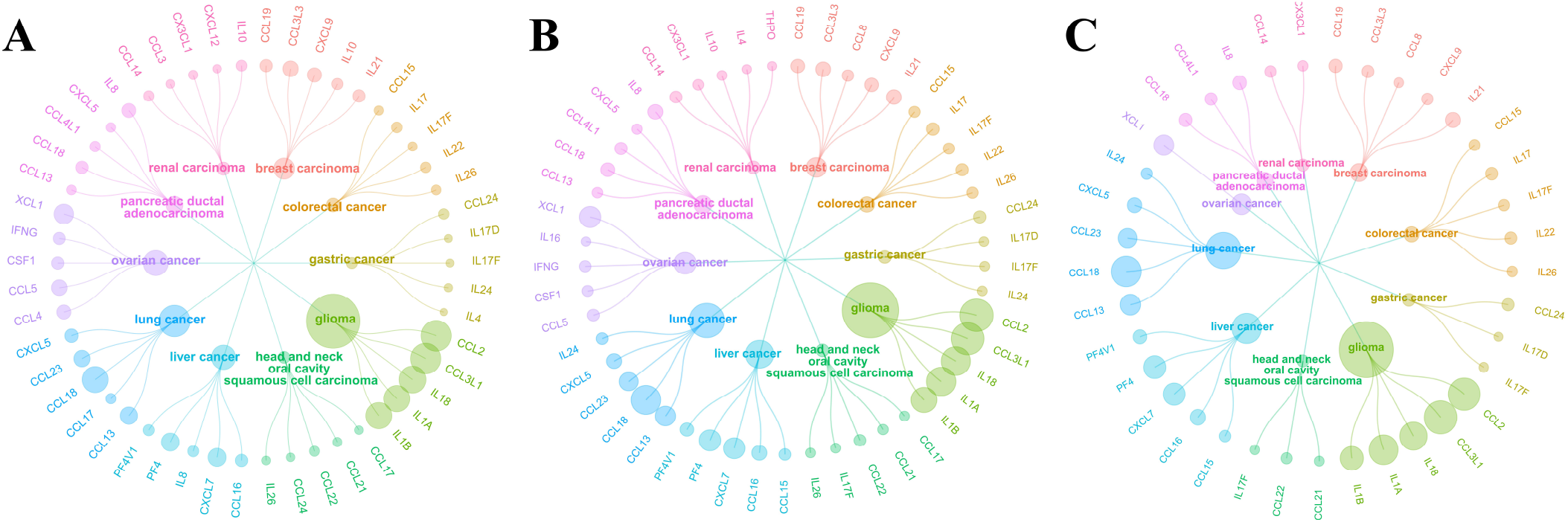
Specific Cytokines in TIME. Figure A, B, and C show the top 5 immune regulatory cytokines identified in the 10 cancers with the highest sample counts under three different threshold values.

## IV. DISCUSSION

Single-cell technology has revolutionized the understanding of the heterogeneity and functional diversity of individual cells, bringing tremendous opportunities to biology and precision medicine. Constructing the single-cell TIME dataset is of great significance for studying applications, such as tumor heterogeneity, immune system, and drug screening. In this study, we devised a custom Python script to effectively gather scRNA-seq data pertaining to the TIME from a diverse range of cancer patients, primarily drawing from public databases, such as GEO. Our method, which integrated Python libraries like BeautifulSoup and requests with parallel techniques using the concurrent futures library, facilitated simultaneous data collection from multiple webpages, significantly expediting the data collection process and resulting in a comprehensive dataset encompassing over 30 cancer types.

Conversely, CancerSCEM [13] and TISCH [27] databases acquire their scRNA-seq data from publicly accessible resources, including GEO, GSA and ArrayExpress, as well as published studies. Although these databases offer valuable resources, the quantity of datasets they incorporated is comparatively limited. Our approach presented more efficient means of data acquisition, enabling the creation of a refined dataset focused exclusively on tumor samples with accompanying information about tissue origin. In terms of data processing, we established an end-to-end pipeline oriented towards TIME, including cell quality control, immune microenvironment screening, clustering, and cell-type annotation. At the cell-type annotation stage, recognition of immune cells is mainly focused in this study. Unlike the complex mode of combined annotation tools and manual operations, we only used SingleR’s recognition algorithm and regard marker genes as hyperparameters. After multiple finetunes, a set of highly differential marker genes is determined which achieved good recognition results on the dataset. The specificity of the data collection and processing strategy empoared us to generate accurate insights into the TIME and its involvement across various cancer types, ultimately contributing to the continued advancement of cancer immunity research.

In recent years, artificial intelligence (AI) techniques such as ML and deep learning have redefined the methods and capabilities for analyzing large-scale data, including scRNA-seq results. Many studies have demonstrated the feasibility of using AI techniques for scRNA-seq data analysis. For example, SAUCIE applied autoencoder to a dataset of 11 million T cells from 180 dengue fever patients, identifying clustering features of acute dengue infection and stratified immune response to dengue fever [28]. Similarly, scGNN combined graph neural network (GNN) with scRNA-seq data to identify 10 neuronal clusters and cell-type-specific markers in Alzheimer’s disease [29]. SpaGCN, based on spatially resolved transcriptomics data, identified cancerous and non-cancerous regions of human primary pancreatic tumors and identified two marker genes that distinguish cancerous regions [30]. Dohmen et al. used ML models to identify tumor cells and normal cells in tumor single-cell data [31]. The present study focused on studying the TIME by combining ML methods with scRNA-seq data. A TIME dataset containing 39 cancers is constructed and random forest is trained to model the relationship between immune regulatory cytokines and cancer types, achieving high accuracy in the 10 cancers with the highest amount of sample (Figure S4). The results clearly demonstrated that regardless of the universally-expressed cytokines in TIME of all cancers, unique cytokine expression patterns could be recognized for each particular cancer type, which might be further developed as treatment targets or diagnostic markers. The most highly expressed cytokines across all cancer types, such as CCL4 (protein: MIP-1*β*) and IL32 (protein: IL-32), have already been evaluated as drug targets for various malignancies, including acute lymphocytic leukemia, breast cancer, colorectal carcinoma and melanoma [32], [33]. In addition, most of these cytokines are known as pro-inflammatory factors, which further confirmed that mediators and cellular effectors of inflammation are essential constituents of TIME in all kinds of cancers [34], providing valuable information for future studies on inflammatory regulations of immune cells in TIME. The identified cytokines can be further attributed to multiple signaling pathways using online tools such as KEGG Mapper [35], which can be utilized for future explorations on the particular immune cells and signaling pathways related to these cancer types. Based on the splendid potential applications of these results generated in this study, we tried to gather as many scRNA-seq samples as possible. However, the study is still limited by data scale and is difficult to achieve optimized performance in more complicated TIME visualization tasks. Follow-up studies will be continuously performed to extend the dataset and apply advanced deep learning models to model the TIME regulations for cancer diagnosis and treatment.

## V. Conclusion

In this paper, we introduce a current maximum scale scRNA-seq dataset of the TIME and pioneer utilize the machine learning method to model the relationship between cytokine expression patterns and cancer types. The experimental results demonstrate that the proposed method is effective for cancer classification on TIME data, achieving a good accuracy even with a limited amount of cytokine expression data. We believe that this work provides a promising foundation for the development of novel cancer immune diagnosis and treatment strategies based on cytokine immunoregulatory patterns.

## Supporting information

Supplemental materials, and will be used to illustrate the results of the manuscript

## Supplementary Materials

Table S1 shows the 39 human cancers including in this study, which is across 695 human tumor tissues of 117 projects; Table S2 shows the customized marker genes for tumor immune microenvironment identification; The delta median of each cell type in Figure S1 quantifies the recognition confidence of SingleR; Figure S2: A. The proportion of different immune cells in the TIME of the 18 cancer (sample size ≥10). B. The expression of the 20 cytokines with the highest average expression in the TIME of the 18 cancer; Figure S3 shows the classification results of random forest, decision tree, and MLP models and compared the efficacy of each model. Figure S4 is utilized to visualize the Table II data, showing the classification effect of models using different combinations of cytokines.

## Acknowledgments

The authors appreciate Yue Zhou from Peng Cheng Laboratory for the technical advice. The research is supported by the Peng Cheng Laboratory and Peng Cheng Cloud-Brain.

## Notes

This work is funded by the National Natural Science Foundation of China [grant No. 42177417]. The project is supported by the Peng Cheng Laboratory and Peng Cheng Cloud-Brain. This work is also funded by Shanghai Frontiers Science Center of Optogenetic Techniques for Cell Metabolism (Shanghai Municipal Education Commission).

https://sandbox.zenodo.org/record/1180710

### Competing Interest Statement

The authors have declared no competing interest.

### Summary of Updates

Modify the name of institute

## Reference

[1] T. F. Gajewski, H. Schreiber, and Y.-X. Fu, “Innate and adaptive immune cells in the tumor microenvironment,” Nature immunology, vol. 14, no. 10, pp. 1014–1022, 2013.

[2] T. Fu, L.-J. Dai, S.-Y. Wu, Y. Xiao, D. Ma, Y.-Z. Jiang, and Z.-M. Shao, “Spatial architecture of the immune microenvironment orchestrates tumor immunity and therapeutic response,” Journal of hematology & oncology, vol. 14, no. 1, p. 98, 2021.

[3] M. Marzagalli, N. D. Ebelt, and E. R. Manuel, “Unraveling the crosstalk between melanoma and immune cells in the tumor microenvironment,” in Seminars in cancer biology, vol. 59. Elsevier, 2019, pp. 236–250.

[4] L. Li, R. Yu, T. Cai, Z. Chen, M. Lan, T. Zou, B. Wang, Q. Wang, Y. Zhao, and Y. Cai, “Effects of immune cells and cytokines on inflammation and immunosuppression in the tumor microenvironment,” International Immunopharmacology, vol. 88, p. 106939, 2020.

[5] C. A. Dinarello, “Historical insights into cytokines,” European journal of immunology, vol. 37, no. S1, pp. S34–S45, 2007.

[6] D. J. Propper and F. R. Balkwill, “Harnessing cytokines and chemokines for cancer therapy,” Nature reviews Clinical oncology, vol. 19, no. 4, pp. 237–253, 2022.

[7] P. Berraondo, M. F. Sanmamed, M. C. Ochoa, I. Etxeberria, M. A. Aznar, J. L. Pérez-Gracia, M. E. Rodríguez-Ruiz, M. Ponz-Sarvise, E. Castañón, and I. Melero, “Cytokines in clinical cancer immunotherapy,” British journal of cancer, vol. 120, no. 1, pp. 6–15, 2019.

[8] B. E. Lippitz, “Cytokine patterns in patients with cancer: a systematic review,” The lancet oncology, vol. 14, no. 6, pp. e218–e228, 2013.

[9] A. Autenshlyus, A. Studenikina, N. Varaksin, and V. Lyakhovich, “Cytokine production by tumor bioptate at different pathological prognostic stages in breast cancer,” in Doklady Biochemistry and Biophysics, vol. 497, no. 1. Springer, 2021, pp. 86–89.

[10] A. Bellesoeur, N. Torossian, S. Amigorena, and E. Romano, “Advances in theranostic biomarkers for tumor immunotherapy,” Current Opinion in Chemical Biology, vol. 56, pp. 79–90, 2020.

[11] C. Alfaro, M. F. Sanmamed, M. E. Rodríguez-Ruiz, Á. Teijeira, C. Oñate, Á. GonzÁlez, M. Ponz, K. A. Schalper, J. L. Pérez-Gracia, and I. Melero, “Interleukin-8 in cancer pathogenesis, treatment and follow-up,” Cancer treatment reviews, vol. 60, pp. 24–31, 2017.

[12] S. Slovin, A. Carissimo, F. Panariello, A. Grimaldi, V. Bouché, G. Gambardella, and D. Cacchiarelli, “Single-cell rna sequencing analysis: a step-by-step overview,” RNA Bioinformatics, pp. 343–365, 2021.

[13] J. Zeng, Y. Zhang, Y. Shang, J. Mai, S. Shi, M. Lu, C. Bu, Z. Zhang, Z. Zhang, Y. Li et al., “Cancerscem: a database of single-cell expression map across various human cancers,” Nucleic acids research, vol. 50, no. D1, pp. D1147–D1155, 2022.

[14] E. Christensen, A. Naidas, D. Chen, M. Husic, and P. Shooshtari, “Tmexplorer: A tumour microenvironment single-cell rnaseq database and search tool,” Plos one, vol. 17, no. 9, p. e0272302, 2022.

[15] B.-H. Kim, K. Yu, and P. C. Lee, “Cancer classification of single-cell gene expression data by neural network,” Bioinformatics, vol. 36, no. 5, pp. 1360–1366, 2020.

[16] D. Mercatelli, F. Ray, and F. M. Giorgi, “Pan-cancer and single-cell modeling of genomic alterations through gene expression,” Frontiers in genetics, vol. 10, p. 671, 2019.

[17] K. F. Mahin, M. Robiuddin, M. Islam, S. Ashraf, F. Yeasmin, and S. Shatabda, “Panclassif: Improving pan cancer classification of single cell rna-seq gene expression data using machine learning,” Genomics, vol. 114, no. 2, p. 110264, 2022.

[18] T. Barrett, S. E. Wilhite, P. Ledoux, C. Evangelista, I. F. Kim, M. Tomashevsky, K. A. Marshall, K. H. Phillippy, P. M. Sherman, M. Holko et al., “Ncbi geo: archive for functional genomics data sets—update,” Nucleic acids research, vol. 41, no. D1, pp. D991–D995, 2012.

[19] A. Athar, A. Füllgrabe, N. George, H. Iqbal, L. Huerta, A. Ali, C. Snow, N. A. Fonseca, R. Petryszak, I. Papatheodorou et al., “Arrayexpress update–from bulk to single-cell expression data,” Nucleic acids research, vol. 47, no. D1, pp. D711–D715, 2019.

[20] I. Papatheodorou, P. Moreno, J. Manning, A. M.-P. Fuentes, N. George, S. Fexova, N. A. Fonseca, A. Füllgrabe, M. Green, N. Huang et al., “Expression atlas update: from tissues to single cells,” Nucleic acids research, vol. 48,no. D1, pp. D77–D83, 2020.

[21] Y. Hao, S. Hao, E. Andersen-Nissen, W. M. Mauck, S. Zheng, A. Butler, M. J. Lee, A. J. Wilk, C. Darby, M. Zager et al., “Integrated analysis of multimodal single-cell data,” Cell, vol. 184, no. 13, pp. 3573–3587, 2021.

[22] D. Aran, A. P. Looney, L. Liu, E. Wu, V. Fong, A. Hsu, S. Chak, R. P. Naikawadi, P. J. Wolters, A. R. Abate et al., “Reference-based analysis of lung single-cell sequencing reveals a transitional profibrotic macrophage,” Nature immunology, vol. 20, no. 2, pp. 163–172, 2019.

[23] N. A. Mabbott, J. K. Baillie, H. Brown, T. C. Freeman, and D. A. Hume, “An expression atlas of human primary cells: inference of gene function from coexpression networks,” BMC genomics, vol. 14, no. 1, pp. 1–13, 2013.

[24] G. Pizzolato, H. Kaminski, M. Tosolini, D.-M. Franchini, F. Pont, F. Martins, C. Valle, D. Labourdette, S. Cadot, A. Quillet-Mary et al., “Single-cell rna sequencing unveils the shared and the distinct cytotoxic hallmarks of human tcrvδ1 and tcrvδ2 γδ t lymphocytes,” Proceedings of the National Academy of Sciences, vol. 116, no. 24, pp. 11 906–11 915, 2019.

[25] C. M.-C. Li, H. Shapiro, C. Tsiobikas, L. M. Selfors, H. Chen, J. Rosenbluth, K. Moore, K. P. Gupta, G. K. Gray, Y. Oren et al., “Aging-associated alterations in mammary epithelia and stroma revealed by single-cell rna sequencing,” Cell reports, vol. 33, no. 13, 2020.

[26] A. Abraham, F. Pedregosa, M. Eickenberg, P. Gervais, A. Mueller, J. Kossaifi, A. Gramfort, B. Thirion, and G. Varoquaux, “Machine learning for neuroimaging with scikit-learn,” Frontiers in neuroinformatics, vol. 8, p. 14, 2014.

[27] D. Sun, J. Wang, Y. Han, X. Dong, J. Ge, R. Zheng, X. Shi, B. Wang, Z. Li, P. Ren et al., “Tisch: a comprehensive web resource enabling interactive single-cell transcriptome visualization of tumor microenvironment,” Nucleic acids research, vol. 49, no. D1, pp. D1420–D1430, 2021.

[28] M. Amodio, D. Van Dijk, K. Srinivasan, W. S. Chen, H. Mohsen, K. R. Moon, A. Campbell, Y. Zhao, X. Wang, M. Venkataswamy et al., “Exploring single-cell data with deep multitasking neural networks,” Nature methods, vol. 16, no. 11, pp. 1139–1145, 2019.

[29] J. Wang, A. Ma, Y. Chang, J. Gong, Y. Jiang, R. Qi, C. Wang, H. Fu, Q. Ma, and D. Xu, “scgnn is a novel graph neural network framework for single-cell rna-seq analyses,” Nature communications, vol. 12, no. 1, p. 1882, 2021.

[30] J. Hu, X. Li, K. Coleman, A. Schroeder, N. Ma, D. J. Irwin, E. B. Lee, R. T. Shinohara, and M. Li, “Spagcn: Integrating gene expression, spatial location and histology to identify spatial domains and spatially variable genes by graph convolutional network,” Nature methods, vol. 18, no. 11, pp. 1342–1351, 2021.

[31] J. Dohmen, A. Baranovskii, J. Ronen, B. Uyar, V. Franke, and A. Akalin, “Identifying tumor cells at the single-cell level using machine learning,” Genome Biology, vol. 23, no. 1, p. 123, 2022.

[32] N. Mukaida, S.-i. Sasaki, and T. Baba, “Ccl4 signaling in the tumor microenvironment,” Tumor Microenvironment: The Role of Chemokines– Part A, pp. 23–32, 2020.

[33] J. T. Hong, D. J. Son, C. K. Lee, D.-Y. Yoon, D. H. Lee, and M. H. Park, “Interleukin 32, inflammation and cancer,” Pharmacology & therapeutics, vol. 174, pp. 127–137, 2017.

[34] C. I. Diakos, K. A. Charles, D. C. McMillan, and S. J. Clarke, “Cancer-related inflammation and treatment effectiveness,” The Lancet Oncology, vol. 15, no. 11, pp. e493–e503, 2014.

[35] M. Kanehisa, M. Furumichi, Y. Sato, M. Kawashima, and M. Ishiguro-Watanabe, “Kegg for taxonomy-based analysis of pathways and genomes,” Nucleic acids research, vol. 51, no. D1, pp. D587–D592, 2023.

