## Supplemental materials, and will be used to illustrate the results of the manuscript for "Cytokine expression patterns: A single-cell RNA sequencing and machine learning based roadmap for cancer classification"

Table of Contents

| **Table S1** | S2 |
| --- | --- |
| **Table S2** | S3 |
| **Figure S1** | S4 |
| **Figure S2** | S5 |
| **Figure S3** | S6 |
| **Figure S4** | S7 |

**Table S1.** 39 human cancers including in this study, which is across 695 human tumor tissues of 117 projects

| **39 Cancer Types Across the Dataset**  **Marker Genes** | |
| --- | --- |
| acute myeloid leukemia  anaplastic thyroid cancer  bladder cancer  breast carcinoma  cavernous hemangioma  cervical cancer  cholangio carcinoma  chronic lymphocytic leukemia  colorectal cancer  cushing's disease adenoma  cutaneous squamous cell carcinoma  endometrioma  ependymoma  esophageal cancer  fallopian tube cancer  gastric cancer  glioma  head and neck oral cavity squamous cell carcinoma  high-grade serous fallopian tube carcinoma  human giant cell tumor | liver cancer  lung cancer  lymphoma  melanoma  meningioma  mesothelioma  metastatic pancreatic neuroendocrine tumor  myeloma  ovarian cancer  pancreatic ductal adenocarcinoma  papillary thyroid carcinoma  pleuropulmonary blastoma  prostate cancer  renal carcinoma  retinoblastoma  sarcoma  sialoblastoma  skin cancer  testicular cancer |

**Table S2. Marker genes for TIME identification.** The table shows a set of customized marker genes for filtering immune cells from tumor tissue.

| **Cell Type** | **Marker Genes** |
| --- | --- |
| B cell  GMP  Neutrophil  NK cell  T cell  Macro/Mono/DC HSC | CD19, MS4A1, BANK1  CD38, KIT, ADK  ADGRG3, CXCL8, FCGR3B  FCGR3A, KLRB1, KLRD1  CD3D, CD3G, CD3E  CD68, CD14, MRC1  CD34, ITGA5, PROM1 |

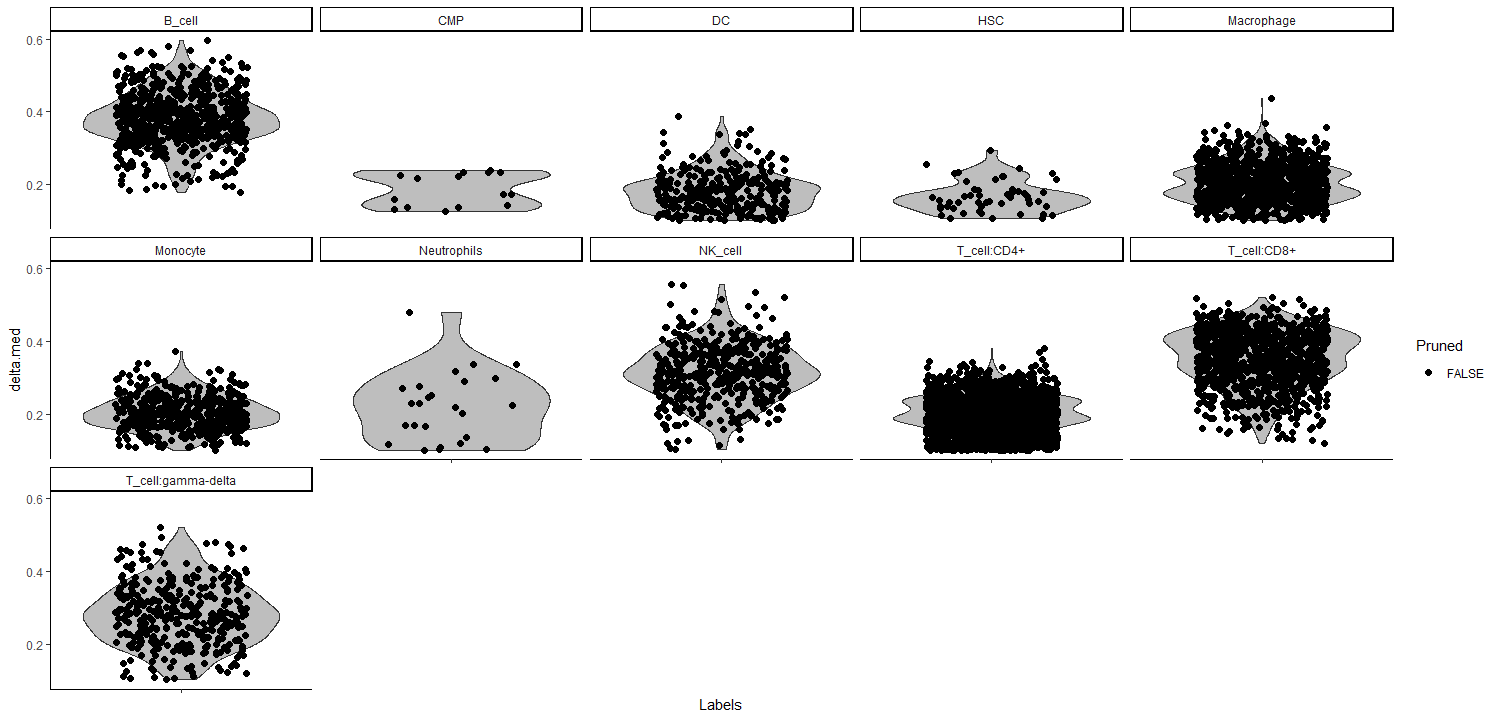

**Figure S1. Delta Median of Cell Types**. The figure quantifies the recognition confidence of SingleR. The minimum acceptable delta was set to 0.1.

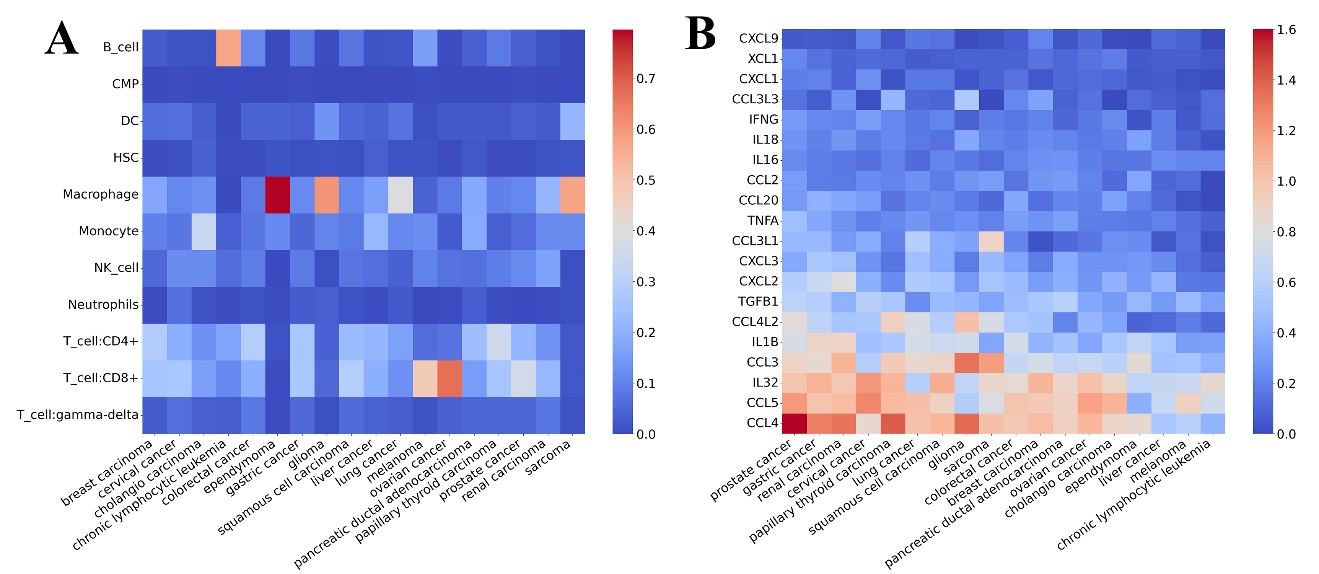

**Figure S2. Statistics of TIME.** A. The proportion of different immune cells in the TIME of the 18 cancer (sample size ≥10). B. The expression of the 20 cytokines with the highest average expression in the TIME of the 18 cancer.

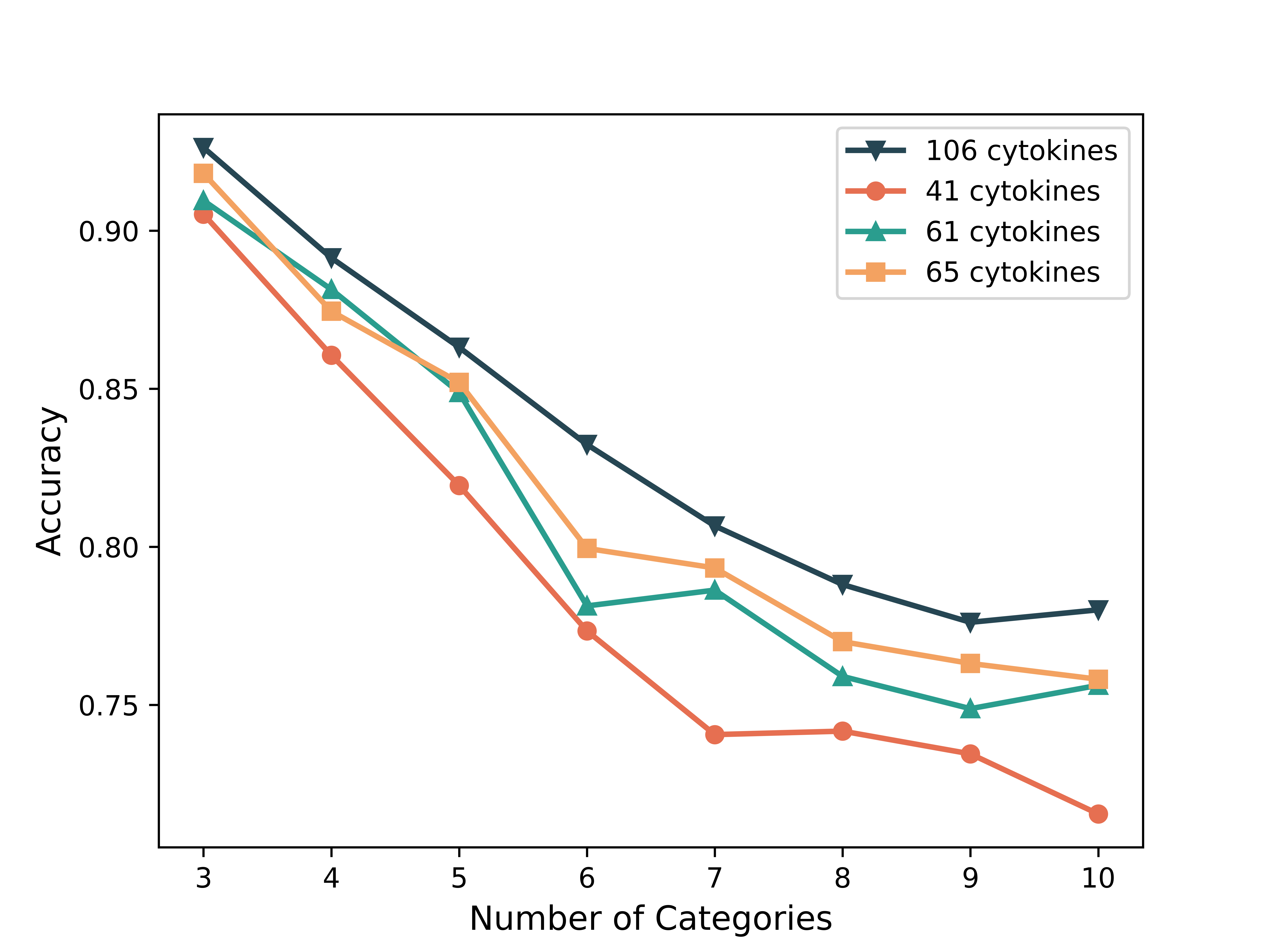

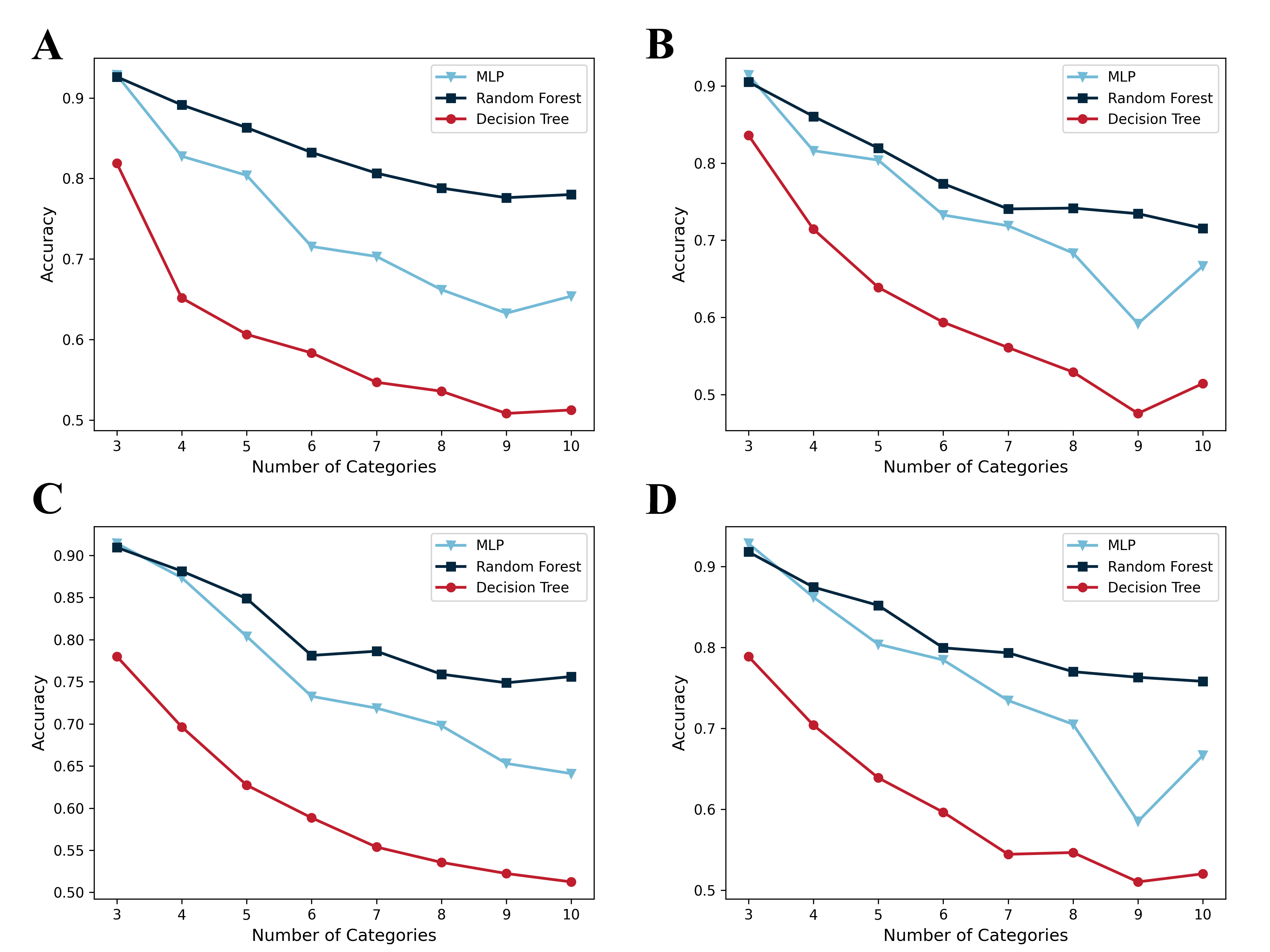
**Figure S3. Classification Results of different models.** The efficacy of random forest, decision tree, and MLP models were compared using different sets of training data. The results, denoted as A, B, C, and D, using data containing 106, 41, 61, and 65 cytokines, respectively.

**Figure S4. Classification Results of Random Forest with Different Cytokines.** Visualization of Table 2 data, showing the classification effect of models using different combinations of cytokines.
